# Evaluation of Fecal Immunoassays for Canine *Echinococcus* Infection in China

**DOI:** 10.1101/2020.08.11.246009

**Authors:** Liying Wang, Qian Wang, Huixia Cai, Hu Wang, Yan Huang, Yu Feng, Xuefei Bai, Min Qin, Sylvie Manguin, Laurent Gavotte, Weiping Wu, Roger Frutos

## Abstract

Human echinococcosis is present worldwide but it is in China that prevalence is the highest. Western China and in particular the Tibetan plateau is the region where the burden of echinococcosis is the most important. Dogs are a major carrier of echinococcosis and monitoring the presence of *Echinococcus* worms in dogs is therefore essential for efficiently controlling the disease. Detection kits based on three different technologies, i.e. sandwich ELISA, ELISA and gold immunodiffusion are currently marketed and used in China. The objective of this work was to assess the efficacy of these kits, in particular with respect to sensitivity and specificity. Four fecal antigen detection kits for canine echinococcosis covering the three technologies were obtained from companies and tested in parallel on 220 fecal samples. The results indicate that the performance is lower than expected, in particular in terms of sensitivity. The best results were obtained with the sandwich ELISA technology. The gold immunofiltration yielded the poorest results. In all cases, further development is needed to improve the performance of these kits, which represent a key element for the control of echinococcosis.

**Author summary:** Although present worldwide, human echinococcosis is at its highest prevalence in Western China and particularly on the Tibetan plateau. Controlling echinococcosis is a national priority and routine monitoring must be established. Dogs are the main carriers of echinococcosis and surveying *Echinococcus* worms in dogs is therefore a key issue. Commercial detection kits are currently in use in China for monitoring the presence of Echinococcosis in dogs. These kits are based on three different technologies, i.e. sandwich ELISA with two monoclonal antibodies, ELISA, and gold immunodiffusion. National survey programs are essential for the control of echinococcosis and it is thus very important to assess the efficacy of these kits, planned to be used for the national survey programs. The work was thus undertaken to assess this efficacy, in particular with respect to sensitivity and specificity. Four fecal antigen detection kits for canine echinococcosis covering the three technologies were obtained from companies and tested in parallel on 220 fecal samples. The performance was lower than expected, in particular for their sensitivity, which ranged from 51.5% to 83.9% with only two samples displaying a worm burden lower than 100. Three out of four kits showed non-specific cross-reactions with other parasites. The best results were obtained with the sandwich ELISA technology, whereas gold immunofiltration yielded the poorest results. However, in all cases, further development is strongly needed to improve the performance of these kits, which represent a key element for the control of echinococcosis.

## 1. Introduction

Echinococcosis is a health-threatening parasitic zoonotic disease caused by the larval stage of *Echinococcus* tapeworms [1]. Cystic echinococcosis (CE) and alveolar echinococcosis (AE) in humans, livestock and small mammals are triggered by the involuntary consumption of *Echinococcus granulosus* and *Echinococcus multilocularis* eggs, respectively, which are excreted in the feces of the definitive hosts, i.e. carnivores. Naturally, the transmission occurs between definitive hosts (primary dogs and foxes) and intermediate hosts (livestock and small mammals), whilst humans are accidental hosts. Human infection can occur through direct contact with the definitive host or indirectly through contamination of food or possibly water with parasite eggs [2]. *Echinococcus granulosus* is distributed worldwide, with only a few areas such as Iceland, Ireland, and Greenland, which are considered free of autochthonous human cases [3,4]*. Echinococcus multilocularis* is confined to the northern hemisphere, but within that range displays a wide distribution [5]. In humans, metacestode infection causes severe disease and possibly death. It also results in economic losses from treatment costs, lost wages and livestock-associated production losses. It has been recognized as one of the world’s public health issues.

Both CE and AE are endemic in the pasture areas of western China, threatening more than 50 million people with a global echinococcosis prevalence of 0.28% in humans, 4.68% in livestock and 4.25% in dogs. The number of patients was estimated to be 166,098 [6]. All provinces (autonomous regions, municipalities and special administrative regions) have recorded cases of echinococcosis. Echinococcosis is indigenous in endemic provinces and imported in non-endemic provinces. Echinococcosis has been listed as a key parasitic disease in China [7]. China is believed to be accountable for 40% of the world CE Disability Adjusted Life Years (DALYs) [8]. A national control project has been implemented in echinococcosis endemic areas in western China since 2006. Dog management and monthly treatment with praziquentel are two major intervention measures implemented to prevent human and livestock infections. Therefore, the detection of *Echinococcus* infections in dogs is a very important indicator to assess control efficacy and risk of disease transmission [9].

Diagnosis and detection of *Echinococcus granulosus* (*sensu lato*) infection in animals is a prerequisite for epidemiological studies and surveillance of echinococcosis in endemic, re-emergent or emergent transmission zones. Testing dog fecal samples by coproantigen ELISA, often combined with mass ultrasound screening programs for human CE, has been the preferred approach for monitoring and surveillance in resource-poor endemic areas and during control schemes [10]. Dogs infection rates are very high and sensitive indicators to assess the risk and burden of echinococcosis and to evaluate the effect of control measures [11].

Currently, 2 sandwich ELISA kits and 1 ELISA test for the detection of coproantigen as well as a gold immunofiltration assay are commercially available in China. In this work, we evaluated the relative performance of these four kits representing three different technologies, in particular sensitivity and specificity, in the detection of *Echinococcus granulosus* infections in dogs in order to provide a reference for practical implementation in control projects.

## 2. Materials and methods

### 2.1. Kits assessed

This study assessed 4 kits currently used for the prevention and control of echinococcosis in China. The kits were randomly coded as A, B, C and D. The information on the kits is provided in Table 1. These kits were provided by Xinjiang Tecon Animal Husbandry Bio-Technology Co., Ltd, Zhuhai S.E.Z. Haitai Biological Pharmaceuticals Co., Ltd. and Shenzhen Combined Biotech Co., Ltd (Table 1). Two kits were sandwich ELISA tests (A and B), one is an ELISA test (D) and one is a Gold Immunofiltration assay (C) (Table 1).

**Table 1.**
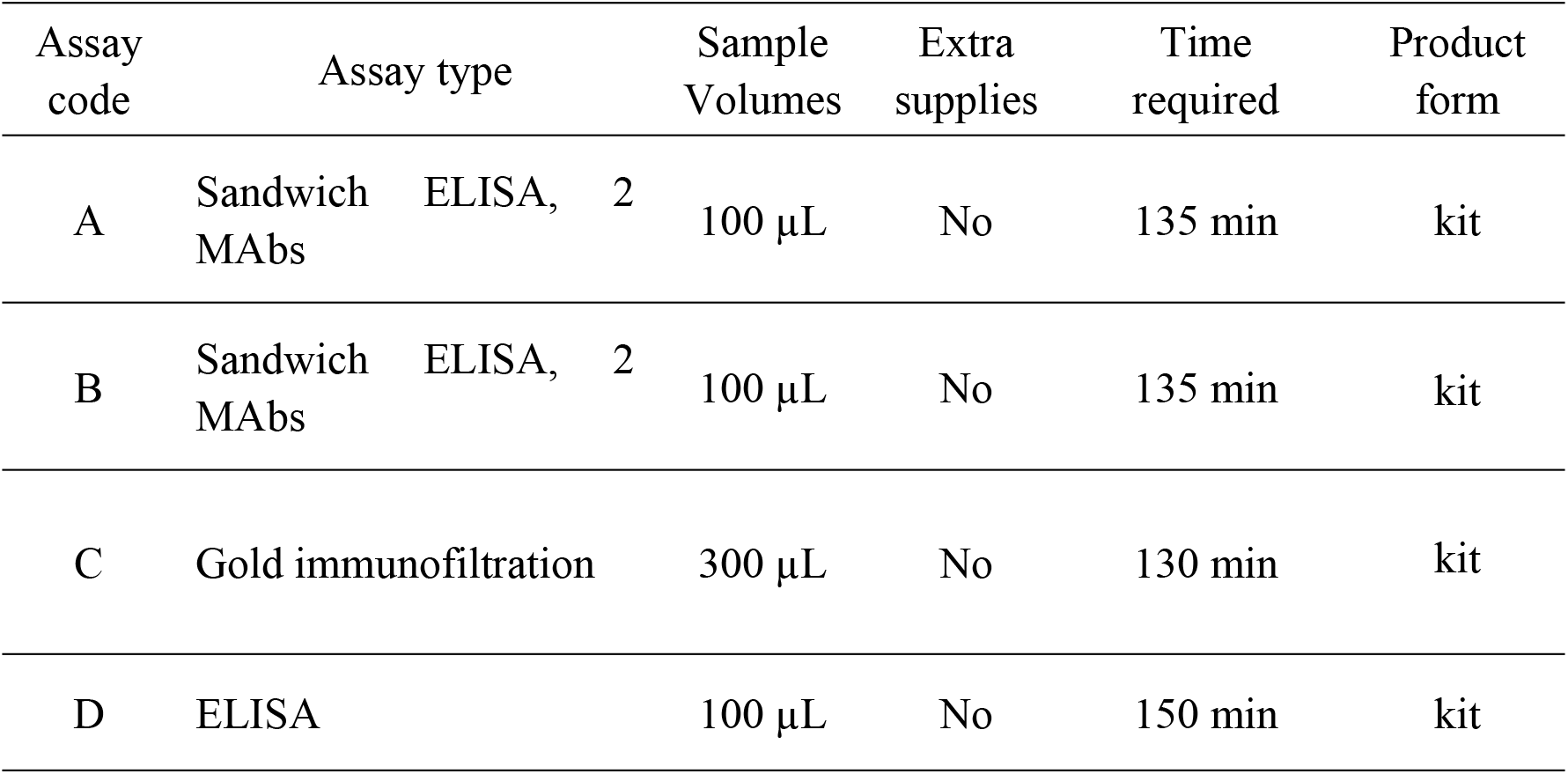
Major features of the assessed tests for *Echinococcus granulosus* diagnosis in dogs.

### 2.2. Specimen collection

A total of 34 positive canine fecal specimens were collected from dogs in Xinjiang, Qinghai and Gansu Positive cases were identified by demonstration of the presence of adult worms in the intestine, which is considered as the “gold” standard for the identification of *Echinococcus* infections [12]. Hence, we detected *E. granulosus* through autopsy in 34 specimens with a minimum parasite load of 5 and a maximum load of 25,000 (Table 2). A complement of 158 negative canine fecal specimens were collected, out of which 116 were from non-endemic areas in Gansu and 42 from laboratory dogs without any parasitic infection. An additional 28 samples of canine fecal specimens were also collected from dogs displaying other parasitic infections. Eight samples of *Taenia hydatigera*, 8 of *Dipylidium caninum* and 12 of *Spirometra mansoni*, were collected in the Guangdong province (Table 3). All specimens were verified by etiologic inspection.

**Table 2.** Specific parasite load in the 34 positive samples.

| Infection Level | Worm load Range | Number of infected dogs | Worm burden |
| --- | --- | --- | --- |
| I | [0-500] | 9 | 5;60;100;200(3);300(2);400 |
| II | [500-5,000] | 10 | 600(2);1,100;1,500;2,100;3,100 (2);3,500(2);4,000 |
| III | [5,000-20,000] | 7 | 6,000(5);11,000(2) |
| IV | [20,000-+∞] | 8 | 20000(3);25000(5) |

**Table 3.** Composition and origin of samples.

| Category | Number of dogs sampled | Sample origin |
| --- | --- | --- |
| Positive canine fecal specimens | 34 (*Echinococcus granulosus* infection) | Qinghai Center for Disease Control and Prevention and The Animal Husbandry Institute of Xinjiang Uygur Autonomous Region |
| Negative canine fecal specimens | 158 (from non-endemic areas in Gansu province) | Gansu Center for Disease Control and Prevention |
| Canine fecal specimens of other parasitic infection | 28 (including 8 with *Taenia hydatigera*, 8 with *Dipylidium caninum*, 12 with *Spirometra mansoni*) | Sun Yat-Sen University of Guangdong province |

### 2.3. Sample preparation

A double-blind method was used in the detection process. All information was kept confidential. Experimenters did not know what they were testing, they only received code numbers as sample identifiers. Each one of the four kits was handled by a different group. In order to ensure that the concentration of sample in the different groups was the same, the preliminary preparation of the samples was performed by the senior experimenter of the organization. Samples were stored at −80°C upon collection. Fecal specimens were defrosted and 3g of each sample were diluted in Phosphate Buffer Saline (PBS) at pH 7.2 ~ 7.4, to the final concentration of 1g/mL and centrifuged at 3000 G for 30 minutes. After centrifugation, 2 mL of supernatant were collected. For two groups of parallel samples for each test, 6 sample batches of 100μL and 2 sample batches of 300μL were prepared. In order to avoid any mutual confirmation of results, all samples were randomly encoded. The information was kept confidential.

### 2.3. Detection tests

All samples were tested by each kit in double according to the manufacturer’s instructions. The operator of each detection kit was assigned by the company. Parallel detection tests with the four different kits were conducted simultaneously in the same laboratory.

### 2.4. Data analysis

Data were analyzed using the SPSS 20.0 software package (IBM, Armonk, USA). The indicators considered for analysis were: accuracy, reliability, sensitivity, specificity, positive predictive value, negative predictive value, Youden’s index, cross reaction rate, consistency rate and weighted consistency rate. Data were tested using a chi-square test. Each index is the average of the test results of two groups of parallel samples. Definitions and calculation methods of relevant indicators are as follows.

#### Sensitivity

Proportion of known infected fecal samples that tested positive in an assay (Infected fecal samples that tested negative are considered as false negatives.)

#### Specificity

Proportion of uninfected reference fecal samples that tested negative in an assay. (Uninfected fecal samples that tested positive are regarded as false positives). This type of specificity is denominated **specificity 1**. Specificity tests which referring to reference fecal samples not infected with *Echinococcus* but harboring other parasites is denominated **specificity 2.**

#### Cross reaction rate

Proportion of samples uninfected with *Echinococcus* but harboring other parasites reference fecal samples, which tested positive in an assay.

#### Positive predictive value (PV+)

PV+ is an indicator of the probability that individuals with positive testing results do have the disease.

#### Negative predictive value (PV−)

The PV− is an indicator of the probability that individuals with negative testing results do not have the disease.

#### Youden’s index

Youden’s index expresses the total ability of a reagent to detect true positive and true negative samples.

#### Consistency rate

Proportion of samples with the same test results of reagents as the real results.

#### Kappa Value

Kappa value is used to analyze and evaluate the consistency of two parallel samples detected by one detection method. Considering the influence of opportunity factors on consistency rate.

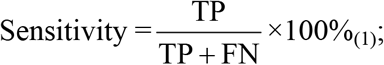

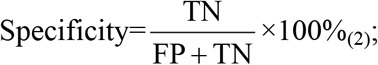

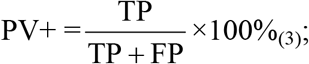

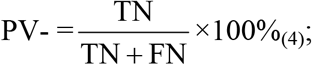

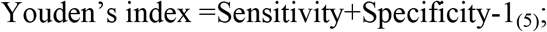

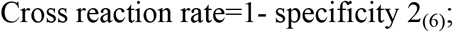

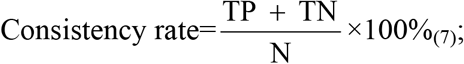

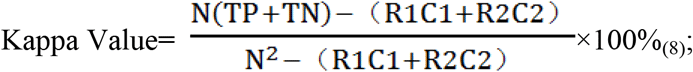

N: total number of samples; TP: true positive; FP: false positive; TN: true negative; FN: false negative; R1: sum of the first row; R2: sum of the second row; C1: sum of the first column; C1: sum of the second column.

Sensibility and specificity normality were confirmed by Kolmogorov Smirnov normality test, thereafter significant differences between assays were assessed by Student’s T-test.

## 3. Results

### 3.1 Sensitivity assessment

The sensitivity of each detection kit was assessed using the 34 feces obtained from *Echinococcus* infected dogs listed in Table 4. We randomly selected 2 parallel groups including 34 fecal specimens and calculated the average as the sensitivity of each detection method. The kit B displayed the highest average sensitivity, i.e. 83.82%, while D showed the lowest average sensitivity, i.e. 51.47% (Tables 4 and 5). The average sensitivity of kits A and C was 76.47% and 70.59%, respectively. When the sensitivity was calculated according to the worm load, strong variations were observed (Table 5). The sensitivity varied widely depending on the worm count. For kit A, the sensibility varied from a lowest of 44.44% for a worm burden class of 500 or less to a maximum of 100% for a worm burden of 5,000 to 20,000. The sensitivity decreased sharply to 81% for a worm burden above 20,000 (Table 5). For the other three kits the calculated sensitivity increased along with the worm load. The lowest sensitivity for a worm load below 500 was 72.22%, 44.44% and 11.11% for kits, B, C and D, respectively (Table 5). The highest sensitivity was observed for a worm burden above 20,000 with 93.75% for kits B and C, and 81.25% for kit A and D (Table 5)

**Table 4.**
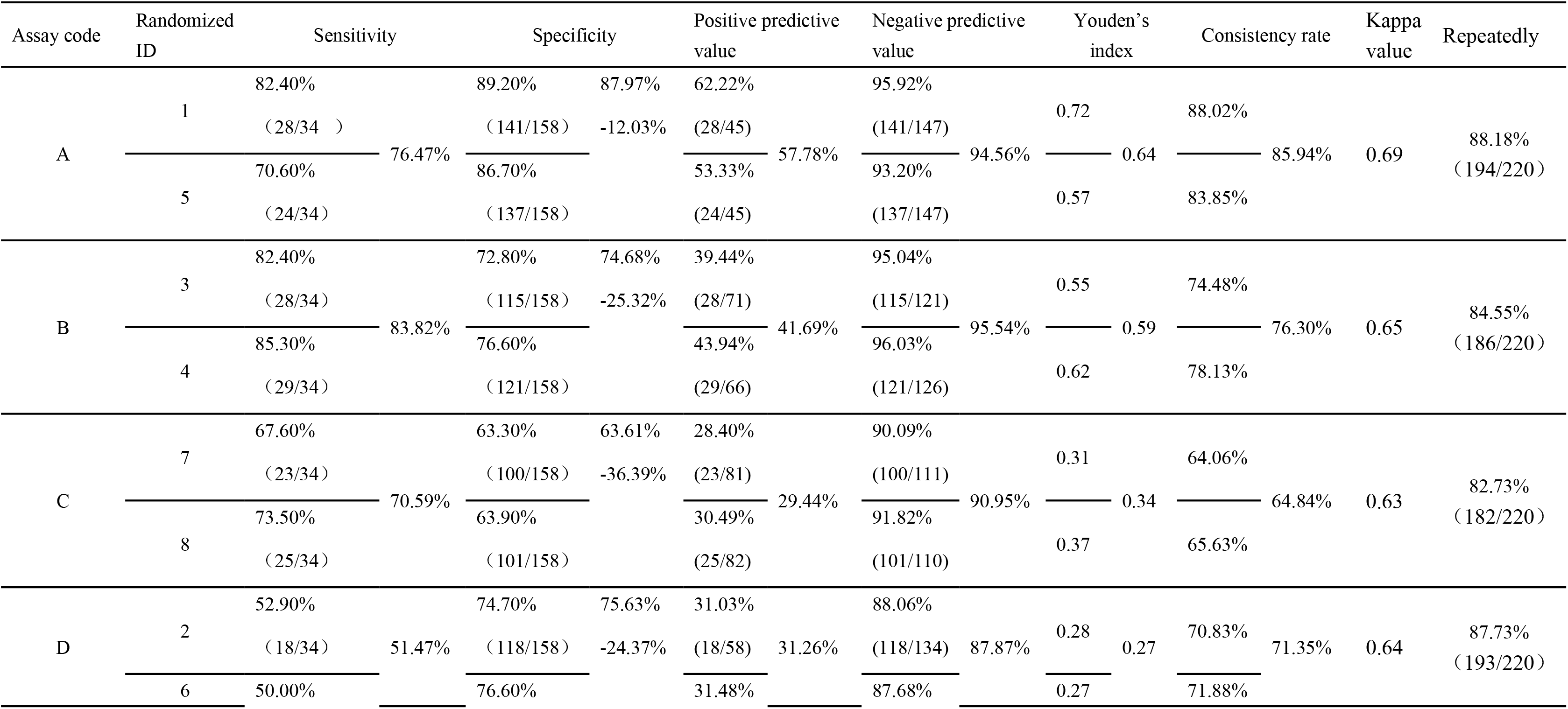
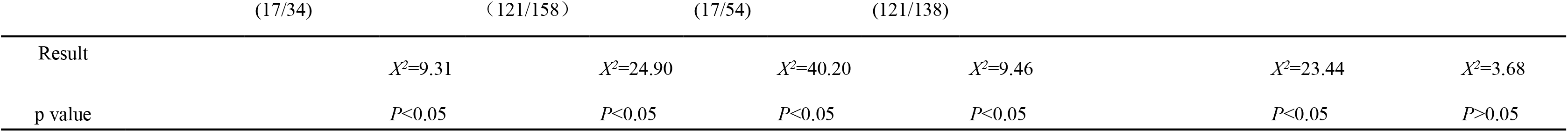
Summary of evaluation results of relevant indicators.

**Table 5.**
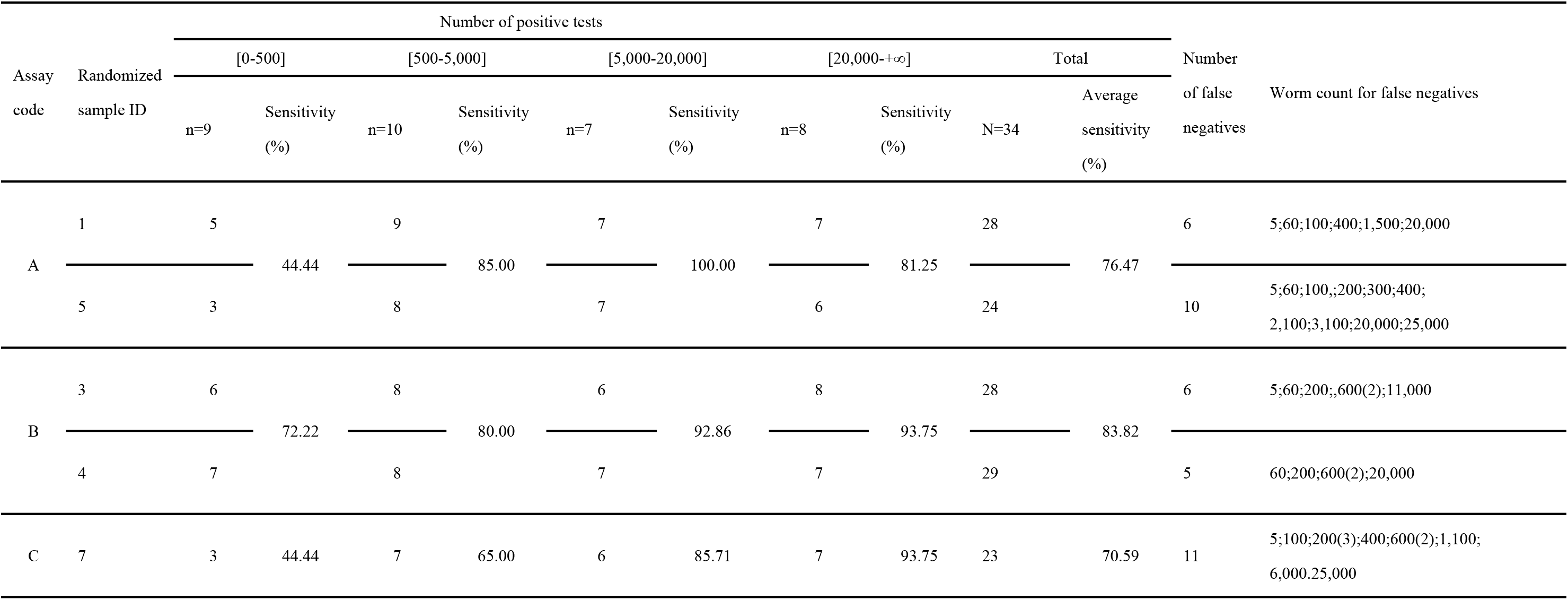
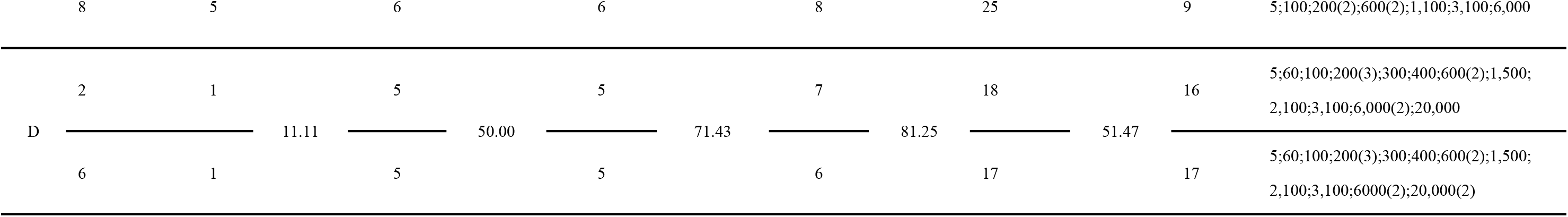
Effect of the worm load sensitivity.

### 3.2. Assessment of false positive answers (specificity 1)

The level of non-specific reactions was assessed for each detection kit on 158 feces obtained from *Echinococcus-*negative dogs (Table 4). We randomly selected 2 parallel groups from the 158 fecal specimens. The lowest level of non-specific reaction was shown by the kit A, i.e. 12.03%, while the kit C displayed the highest level of non-specificity, i.e. 36.39%. Kits B and D yielded intermediate values, i.e. 25.32% and 24.37%, respectively (Table 4).

### 3.3. Cross-reactivity assessment with other tapeworms (specificity 2)

Kit A displayed no cross-reactivity at all with any of the control parasites, i.e. *T. hydatigena*, *D. caninum*, and *S. mansoni* (Table 6). Kits B and C displayed the highest level of cross-reactivity, i.e. 23.21%, whereas kit D showed an intermediate level of 16.07%. Kit C cross-reacted with all three heterologous worms. Kit B showed cross-reactivity with *D. caninum* and *S. mansoni*, while kit D cross-reacted with *T. hydatigena* and *D. caninum*.

**Table 6.** Cross-reactivity with other parasites.

| Assay code | Randomized<br>sample ID | Number of positive tests |  |  |  | Cross reaction rate<br>(%) |
| --- | --- | --- | --- | --- | --- | --- |
|  |  | *Taenia*<br>*hydatigena*<br>n=8 | *Dipylidium caninum*<br>n=8 | *Spirometra*<br>*mansoni*<br>n=12 | Total<br>N=28 |  |
| A | 1 | 0 | 0 | 0 | 0 | 0.00 |
|  | 5 | 0 | 0 | 0 | 0 |  |
| B | 3 | 0 | 3 | 3 | 6 | 23.21 |
|  | 4 | 0 | 4 | 3 | 7 |  |
| C | 7 | 3 | 2 | 2 | 7 | 23.21 |
|  | 8 | 6 | 0 | 0 | 6 |  |
| D | 2 | 1 | 5 | 0 | 6 | 16.07 |
|  | 6 | 1 | 2 | 0 | 3 |  |

### 3.4. Assessment of global performance and accuracy

The best score when using Youden’s index was obtained by kit A (0.64), whereas kit B reached a score of 0.59 (Table 4). Kits C and D obtained very low scores of 0.34 and 0.27, respectively (Table 4). The Youden’s index varies from 0 to 1 with 0 indicating an undiscriminating, therefore useless test, while 1 is indicating a perfect test. Even with the best scores, kits A and B were far from being perfect. The accuracy assessment conducted to evaluate the repeatability of each test yielded scores higher than 80% whatever the kit considered. However, kits A and D reached a higher score, i.e. 88.18% and 87.73%, respectively, compared to kits B and C, i.e. 84.55% and 82.73%, respectively.

### 3.5. Assessment of differences between assays

The relative difference between assays was assessed using a Student’s T-test following the positive normality assessment of the data sensitivity and specificity. The difference between kits was also assessed with a student’s test. For the sensitivity, results from kit A are significantly different from those obtained with kits C and D but not with those from kit B (Table 7). Results from kit B are significantly different from those from kit C and kit D (Table 7). Results from kit C are not significantly different from those obtained with Kit D (Table 7). With respect to specificity, results from kit A are not significantly different from those coming from kits B and C but are significantly different from those obtained with kit D (Table 8). Kit B yielded results significantly different from those from kits C and D (Table 8). Finally, results from kits C and D were not significantly different (Table 8).

**Table 7.** Assessment of the results difference for sensitivity.

| Comparisons | DoF | T | p-value |
| --- | --- | --- | --- |
| A-B | 2 | 3.23 | 0.084 |
| A-C | 2 | 6.21 | **0.025\*** |
| A-D | 2 | 5.96 | **0.027\*** |
| B-C | 2 | 4.93 | **0.038\*** |
| B-D | 2 | 4.61 | **0.044\*** |
| C-D | 2 | -1.70 | 0.232 |

\* Significant at 0.05
DoF: Degree of Freedom

**Table 8.** Assessment of the results difference for specificity.

| Comparisons | DoF | t | p-value |
| --- | --- | --- | --- |
| A-B | 2 | -0.68 | 0.569 |
| A-C | 2 | 2.24 | 0.155 |
| A-D | 2 | 4.87 | **0.040\*** |
| B-C | 2 | 4.60 | **0.044\*** |
| B-D | 2 | 14.45 | **0.005\*\*\*** |
| C-D | 2 | 3.49 | 0.073 |

\* Significant at 0.05
\*\*\* Significant at 0.01
DoF: Degree of Freedom

## 3. Discussion

China is reporting the highest human prevalence rate of echinococcosis worldwide. Following the 2012-2016 national survey, 368 out of 413 counties were identified as endemic for echinococcosis [6]. Currently the number of endemic counties rose to 370, after the disease was detected in 2017 in Dongxiang County in Gansu province and in the Ulagai Management District in Inner Mongolia (Figure 1). The endemic counties spread over 9 provinces or autonomous regions of Tibet, Sichuan, Qinghai, Xinjiang, Gansu, Ningxia, Inner Mongolia, Yunnan and Shanxi (Figure 1). The overall detection rate was 0.51% (5,133/1,001,173) [6]. The prevalence was estimated to be 0.28% in endemic areas and the number of patients was estimated to be 166,098 with a number of persons at risk of about 60 million [6]. Out of the 370 endemic counties, 158 are located in Qinghai Tibetan Plateau with a prevalence of 1.28% which is 10 times higher than the prevalence in non-Qinghai Tibetan Plateau areas, i.e. 0.13% [6].

**Figure 1.**
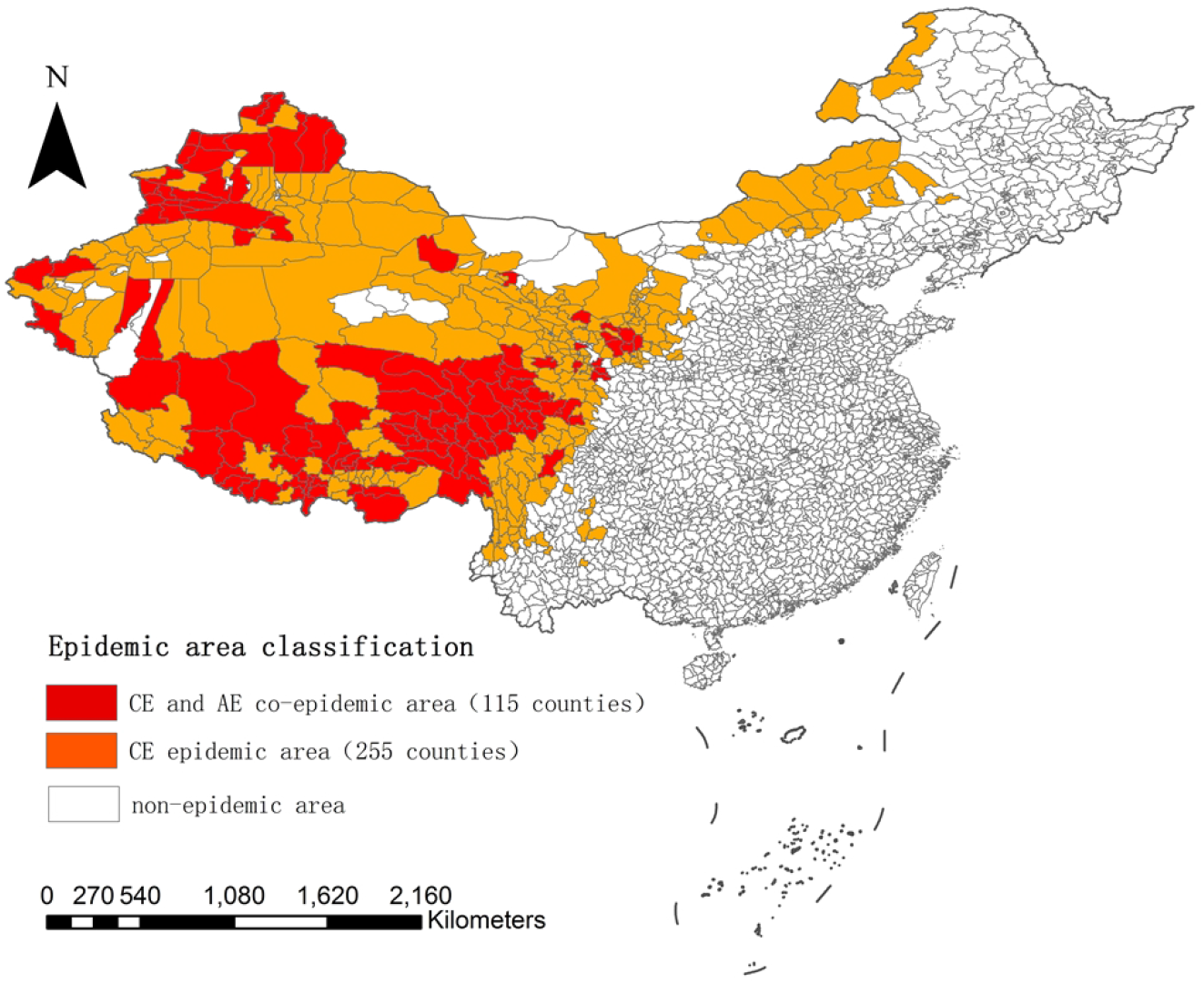
Distribution of echinococcosis by county level in China. AE: Alveolar echinococcosis CE: Cystic echinococcosis

These last few years, significant technical progress was made in immunological diagnosis of *Echinococcus* infection in definitive hosts [13–15]. The detection of parasite antigens in stool by ELISA has been developed for the detection of fecal antigen released by canine *Echinococcus* [13,16–18]. *Echinococcus* antigens can be detected in dog feces 5 to 10 days after being experimentally infected [19]. This detection becomes negative five days after treatment with praziquantel [19]. The detection of specific antigens in the definitive host stool samples is more informative than the detection of serum antibodies, because of the higher probability of being associated with the current infection. Owing to their effectiveness, these tests have been introduced to local echinococcosis prevention programs where they are currently being implemented. ELISA has been adopted as the main diagnostic method in place of the arecoline cathartic method to monitor canine *Echinococcus* infection in control programs. There is thus a need to evaluate the fecal antigen tests regularly to improve the quality of monitoring activities and objectively assess prevention effectiveness.

In this work, we evaluated the accuracy and reliability of four commercial kits currently in use in China and results showed that the sensitivity of the four kits ranged between 51.5% and 83.9% only. Out of all the samples tested, only two displayed a worm burden lower than 100. Therefore, the range of sensitivity obtained in this study is far below that of 92% to 100% for more than 100 worms reported by previous studies for fecal antigen detection [13]. A sensitivity ranging from 29% to 79% has been previously reported for a worm burden lower than 100 worms as determined by necropsy or arecoline cathartic [18]. This is more in the range of what was observed in this work but with a worm burden higher than 100. The sensitivity results reported in this work also indicated that the threshold of 100 is not realistic, which could explain the variation in results from one report to another. The minimal burden of worms for assessing sensitivity should be 500. Nevertheless, owing to the quite low sensitivity observed and to the important variation induced by the worm burden, further modifications and optimization must be conducted to increase the sensitivity of the kits we tested.

Non-specific, or false-positive, reactions from the kits tested in our study ranged from 12.03% to 36.39%, while the cross-reactivity with other parasites ranged from 0 to 23.21%, depending upon the kit. Youden’s index is a comprehensive indicator reflecting the sensitivity and specificity. Under the assumption that sensitivity and specificity are equally important, the kit with the highest Youden’s index is given priority. In this study, the highest Youden’s index was 0.64 for kit A. However, the positive predictive value changes with the infection rate. In this study, the detection results corresponding to the infection rate of 17.70% (34/192) were generally low, indicating the occurrence of false positive results. The consistency rate is the main index reflecting the reliability of kits, which mainly represents the stability of the detection ability of kits. The highest consistency rate of kit A is 85.94%. Kappa value is also an important index to reflect the repeatability of test results. Thus kit A shows the best reliability in terms of repeatability and stability. Although quite high, there is still a need for further improvement in particular because of the level of false positives.

Out of the three technologies assessed, i.e. Sandwich ELISA, ELISA and immunofiltration, the latter displayed the lowest performance score. Immunofiltration has the advantage of being used *in situ* with a simple protocol and with results being immediately available. However, the poor performance displayed by this technology does not make it a reliable and efficient choice for the monitoring of echinococcosis. More developments are needed to improve this technology. This study shows clearly that sandwich ELISA should be the methodology to implement for the surveillance of canine echinococcosis. Kit A displayed the best weighted overall score but kit B yielded a weighted overall score close to the former. Both were based on the technology of sandwich ELISA with two monoclonal antibodies.

ELISA showed performances intermediate between sandwich ELISA and immunofiltration and does not appear as a reliable option for surveillance. Nevertheless, even if sandwich ELISA seems to be the technology of choice for the surveillance of canine echinococcosis, improvements and optimization are still needed to ensure proper surveillance. This study provides a reference for improving control measures and assessment of the prevalence of echinococcosis in the endemic counties of China. This is a key step towards elimination. Since the most sensitive indicator of the risk of epidemic is the dog infection rate, these kits are tools of primary importance. Therefore, we urge manufacturers to strengthen research on their products in order to improve and enhance their overall quality, and in particular sensitivity and specificity, for effective *Echinococcus* diagnosis and control implementation in China.

## Acknowledgements

The authors are very grateful to Xinyu Duan from the first affiliated hospital of Xinjiang Medical University, Xuchu Hu from Sun yat-sen University Guangzhou Province, Zhuangzhi Zhang from the Animal Husbandry Institute of Xinjiang Uygur Autonomous Region, Yu Feng from Gansu Provincial Center for Disease Control and Prevention, Lanzhou, China. and Xiumin Han from Qinghai Provincial People’s Hospital, for their kind help in providing samples.

## Authors participation

Conceived the study: Weiping Wu, Liying Wang, Hu Wang, Qian Wang

Designed the study: Liying Wang

Coordination of units: Weiping Wu

Collected samples and information: Liying Wang

Performed pre-experiment: Huixia Cai

Organization of the work: Liying Wang, Weiping Wu

Supervision: Qian Wang, Hu Wang, Yan Huang, Yu Feng

Data analysis: Liying Wang, Min Qin, Laurent Gavotte, Roger Frutos

Writing-original draft: Liying Wang, Xuefei Bai, Min Qin

Writing-reviewing and editing: All Sylvie Manguin, Laurent Gavotte, Roger Frutos

All authors read and approved the final manuscript.

## References

1. Vuitton DA, McManus DP, Rogan MT, Romig T, Gottstein B, Naidich A, et al. International consensus on terminology to be used in the field of echinococcoses. Parasite. 2020;27:41.

2. Wen H, Vuitton L, Tuxun T, Li J, Vuitton DA, Zhang W, et al. Echinococcosis: Advances in the 21st century. Clin Microbiol Rev. 2019;32:1–39.

3. Craig PS, Rogan MT, Allan JC. Detection, screening and community epidemiology of taeniid cestode zoonoses: cystic echinococcosis, alveolar echinococcosis and neurocysticercosis. Adv Parasitol. 1996;38:169–250.

4. Eckert J, Thompson RC. Historical Aspects of Echinococcosis. Adv Parasitol. 2017;95:1–64.

5. Deplazes P, Rinaldi L, Alvarez Rojas CA, Torgerson PR, Harandi MF, Romig T, et al. Global Distribution of Alveolar and Cystic Echinococcosis. Adv Parasitol. 2017;95:315–493.

6. Wu WP, Wang H, Wang Q, Zhou XN, Wang LY, Zheng CJ, et al. A nationwide sampling survey on echinococcosis in China during 2012-2016, Chin J Parasitol Parasit. 2018,36:1–14. (in chinese) http://www.jsczz.cn/CN/Y2018/V36/I1/1

7. Li B, Quzhen G, Xue CZ, Han S, Chen WQ, Yan XL, et al. Epidemiological survey of echinococcosis in Tibet Autonomous Region of China. Infect Dis Poverty. 2019;8:29.

8. Fasihi Harandi M, Budke CM, Rostami S. The monetary burden of cystic echinococcosis in Iran. PLoS Negl Trop Dis. 2012;6:e1915.

9. Craig PS, Giraudoux P, Wang ZH, Wang Q. Echinococcosis transmission on the Tibetan Plateau. Adv Parasitol. 2019;104:165–246.

10. Craig P, Mastin A, van Kesteren F, Boufana B. *Echinococcus granulosus*: Epidemiology and state-of-the-art of diagnostics in animals. Vet Parasitol. 2015;213:132–148.

11. Craig PS, Hegglin D, Lightowlers MW, Torgerson PR, Wang Q. Echinococcosis: Control and Prevention. Adv Parasitol. 2017;96:55–158.

12. Otero-Abad B, Torgerson PR. A systematic review of the epidemiology of echinococcosis in domestic and wild animals. PLoS Negl Trop Dis. 2013;7:e2249.

13. Zoljargal P, Ganzorig S, Nonaka N, Oku Y, Kamiya M. A survey of canine echinococcosis in Gobi Altai Province of Mongolia by coproantigen detection. Jpn J Vet Res. 2001;49(2):125–129.

14. Nonaka N, Oka M, Kamiya M, Oku Y. A latex agglutination test for the detection of *Echinococcus multilocularis* coproantigen in the definitive hosts. Vet Parasitol. 2008;152:278–283.

15. Machnicka B, Dziemian E, Rocki B, Kołodziej-Sobocińska M. Detection of *Echinococcus multilocularis* antigens in faeces by ELISA. Parasitol Res. 2003;91:491–496.

16. Ahmad G, Nizami WA. Coproantigens: early detection and suitability of an immunodiagnostic method for echinococcosis in dogs. Vet Parasitol. 1998;77:237–244.

17. Allan JC, Craig PS, Garcia Noval J, Mencos F, Liu D, Wang Y, Wen H, et al. Coproantigen detection for immunodiagnosis of echinococcosis and taeniasis in dogs and humans. Parasitology. 1992;104 :347–356.

18. Craig PS, Gasser RB, Parada L, Cabrera P, Parietti S, Borgues C, et al. Diagnosis of canine echinococcosis: comparison of coproantigen and serum antibody tests with arecoline purgation in Uruguay. Vet Parasitol. 1995;56:293–301.

19. Deplazes P, Gottstein B, Eckert J, Jenkins DJ, Ewald D, Jimenez-Palacios S. Detection of *Echinococcus* coproantigens by enzyme-linked immunosorbent assay in dogs, dingoes and foxes. Parasitol Res. 1992;78:303–308.

